# Exploring the energy landscape of bacterial chromosome segregation

**DOI:** 10.1101/2024.07.27.604869

**Authors:** Sumitabha Brahmachari, Antonio B. Oliveira, Matheus F. Mello, Vinícius G. Contessoto, José N. Onuchic

## Abstract

Faithful chromosome segregation during bacterial replication requires global reorganization of the nucleoid, where Structural Maintenance of Chromosomes (SMC) complexes play a crucial role. Here we develop an energy-landscape framework that integrates data-driven pairwise interactions with coarse-grained polymer physics to infer the 3D architectural ensembles of *Escherichia coli* and *Bacillus subtilis* chromosomes throughout replication. We show that SMC-mediated long-range lengthwise compaction reshapes the nucleoid to induce a robust mid-replication transition in which the terminus relocates toward the nucleoid center and duplicated origins segregate toward opposite cell halves. SMC-deficient mutants lack this transition and instead exhibit emergent nematic-like alignment of sister chromosomes that impedes segregation. A distinctive inter-sister Hi-C signature accompanies the emergence of the nematic alignment. By systematically tuning nonspecific inter-sister adhesion, we reveal that SMC activity expands the physical regime permitting faithful segregation. This buffering protects segregation against adhesive forces intrinsic to the crowded bacterial nucleoid. Our framework provides mechanistic insight into SMC-dependent co-replication segregation across bacterial species, yielding experimentally testable predictions for imaging and sister-chromosome-resolved Hi-C.

---

The bacterial chromosome is a densely packed, hierarchically organized polymer whose physical constraints resemble those of eukaryotic genomes, underscoring the universal role of polymer mechanics in chromosome maintenance and reorganization. Structural Maintenance of Chromosomes (SMC) proteins, conserved across all domains of life, are central to this organization: they extrude loops to reinforce long-range intra-chromosome interactions [1, 2]. Steady-state SMC activity via loop extrusion modulates the lengthwise mode of chromosome compaction, facilitating segregation and regulating inter-chromosomal organization across species [3–6]. Length-wise compaction refers to an ordered form of compaction in which loci adjacent along the chromosome contour are brought into close spatial proximity. In bacteria, DNA replication and segregation occur simultaneously. During mid-replication, SMCs interact with nucleoid-associated proteins (NAP) to direct the newly replicated origins (ori) to opposite cell halves, ensuring their segregation [7]. However, a quantitative evaluation of the SMC-driven structural intermediates involved in co-replication segregation of bacterial chromosomes, further consolidating the conserved role of SMC proteins, is lacking.

To dissect how chromosome architecture governs this process, we developed quantitative energy-landscape models for the rod-shaped bacteria *B. subtilis* and *E. coli*. Using a maximum-entropy framework [8–11], we optimized coarse-grained (10 kb) energy landscapes to reproduce Hi-C-derived contact probabilities [12, 13], then extended these landscapes across discrete replication stages to predict the associated three-dimensional structural ensembles. Hi-C-trained models validate emergent features of imaging experiments, such as the axial localizations of ori and ter [14–16]. We performed this analysis for wild-type (`wt`) cells and for SMC mutants, Δ`mukb` lacking the loop-extrusion activity of MukBEF in *E. coli* [13] and Δ`smc`, a mutant of the extrusion complex SMC-ScpAB in *B. subtilis* [12]. Our models predict that SMC-driven long-range lengthwise compaction produces a robust mid-replication transition in which the unreplicated terminus (ter) moves axially inward while replicated origins (ori) segregate outward. This transition, a signature of faithful chromosome segregation, is absent in SMC mutants; they instead show more inter-sister contacts characterized by a nematic alignment of sister chromosome arms that hinder segregation. Across both species, these findings point to a conserved role for SMC proteins in making co-replication segregation more reliable.

Extruding and maintaining stable loops has emerged as the dominant mode of SMC activity. Kinetic modeling of extrusion dynamics has established a paradigm where SMC-mediated loops regulate the steady-state contact probability at loop length scales of ∼ 0.1 − 1 Mb [17–22]. By inferring interaction landscapes directly from data, our framework effectively models steady-state loop extrusion by SMC complexes, together with the ensemble of NAP-driven contacts, enabling a quantitative comparison of their effects on segregation mechanics.

Prior models of bacterial segregation have focused mainly on the entropic effects arising from excluded-volume interactions between self-avoiding polymers [23], with newer work incorporating loop extrusion to direct these entropic forces [24–26]. However, these approaches neglect the heterogeneous adhesive interactions mediated by NAPs and crowders, which also shape global chromosome architecture. Hi-C-constrained polymer reconstructions have shown that experimentally derived contact heterogeneity is essential to recover realistic chromosome architectures in bacteria [27, 28]. Our results, consistent with the previous entropic segregation theories and Hi-C data, show that SMC activity reshapes the chromosome energy landscape to favor entropic segregation forces across species. SMC activity restructures the sister chromosomes to inhibit a NAP-assisted low-entropy nematic state that resists their segregation. The model makes testable predictions, such as an X-shaped Hi-C signature for inter-sister interactions associated with the segregation-deficient nematic state, and provides a conceptual framework to understand the inter-linked roles of NAPs, transcription, and SMCs in coordinating sister-chromosome segregation across bacterial species.

In what follows, we first introduce the essential aspects of the model, learning single-chromosome energy landscapes from Hi-C and constructing the landscapes for partly replicated sister chromosomes (Fig. 1). Then, we inspect the learned single chromosome structures for *E. coli* and *B. subtilis* (Fig. 2), and describe the predictions of the model related to chromosome structure and segregation mechanics using typical inferred parameters (Fig. 3). Subsequently, by varying the parameters and comparing across both species, we link SMC activity to faithful chromosome segregation across a broad physiological context (Fig. 4). Finally, we conclude with a discussion.

**FIG. 1.**
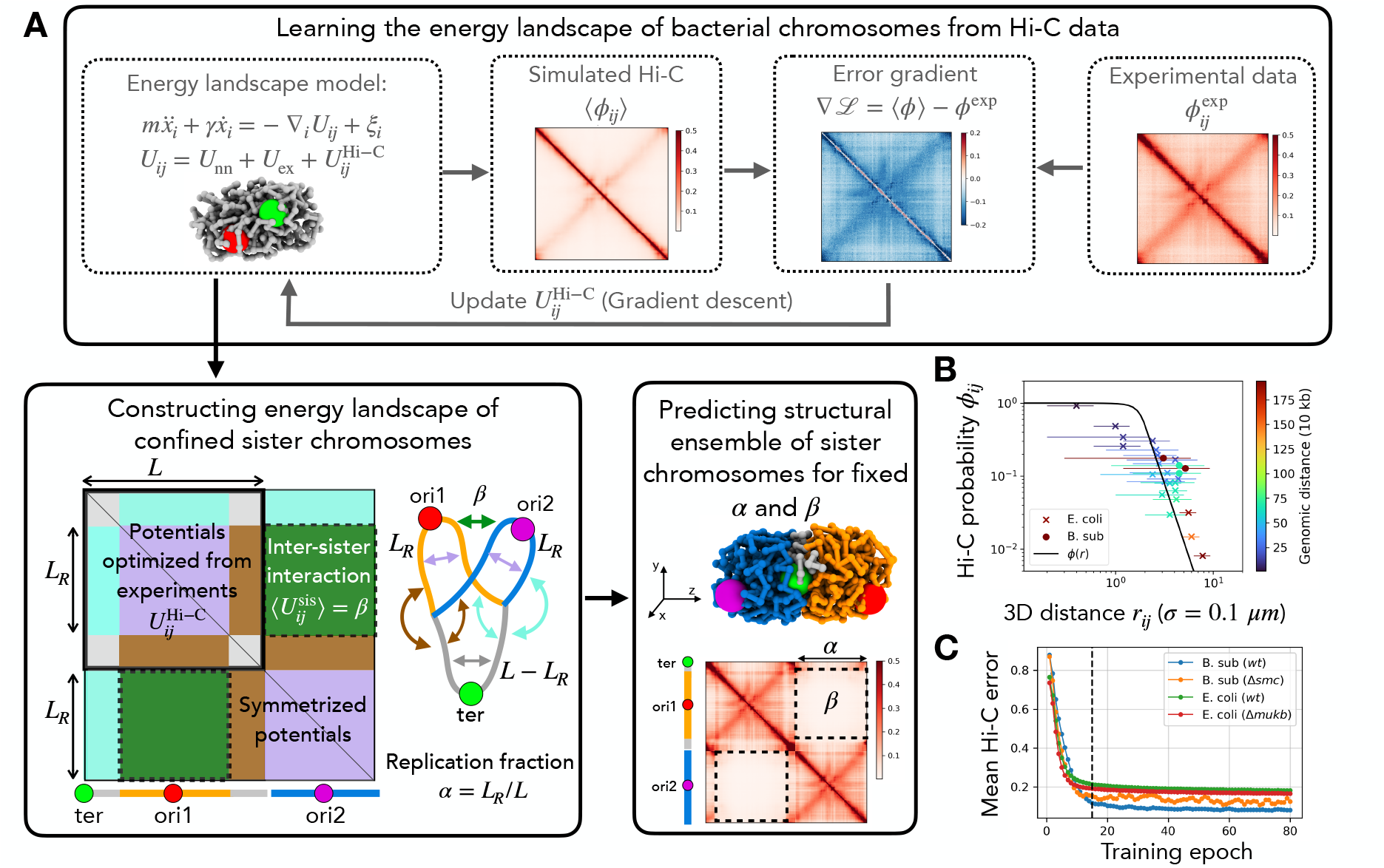
Energy landscape model for replicated sister chromosomes. (A) Schematic of the model construction. The chromosome model consists of a polymer system defined by an energy landscape *U* (*r*), which comprises multiple physics-derived components and one data-driven component 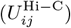. Our goal is to optimize the data-driven term iteratively using a loss function that maximizes entropy within the constraints of the experimental Hi-C data. In each epoch, we compute the pairwise contact probabilities using the simulation model (⟨*ϕ*_*ij*_⟩) and compare them with the observed probabilities *ϕ*^exp^ to backpropagate the error. Once the error converges, we use the learned potential to construct a symmetric energy landscape of sister chromosomes at a certain replication stage, as defined by the replication fraction *α*. The energies corresponding to the inter-sister interactions 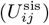 cannot be directly inferred from experimental Hi-C due to sequence similarity of the sisters. We set the mean 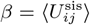 to the non-specific baseline interactions in *U* ^Hi−C^ to study typical behavior, and later sweep this parameter to construct regime diagrams. The constructed model is then used to predict the structural ensembles of partly replicated chromosomes for a given *β* and *α*. (B) Three-dimensional distance between certain loci pairs is plotted against their Hi-C contact probabilities. The markers (diamonds and crosses) represent experimental data where distances are measured using FISH probes [12, 13, 29]; the colors represent the genomic distance between the locus pairs (see colorbar). The solid line is the definition of a Hi-C contact used in the simulations. The horizontal axis is plotted in units of *σ* for the Hi-C function *ϕ* and in units of 0.1 *µ*m for the experimental data. (C) Error convergence during the optimization scheme for various cases. We used the energy landscape at epoch 15 (dashed vertical line) for all downstream analyses.

**FIG. 2.**
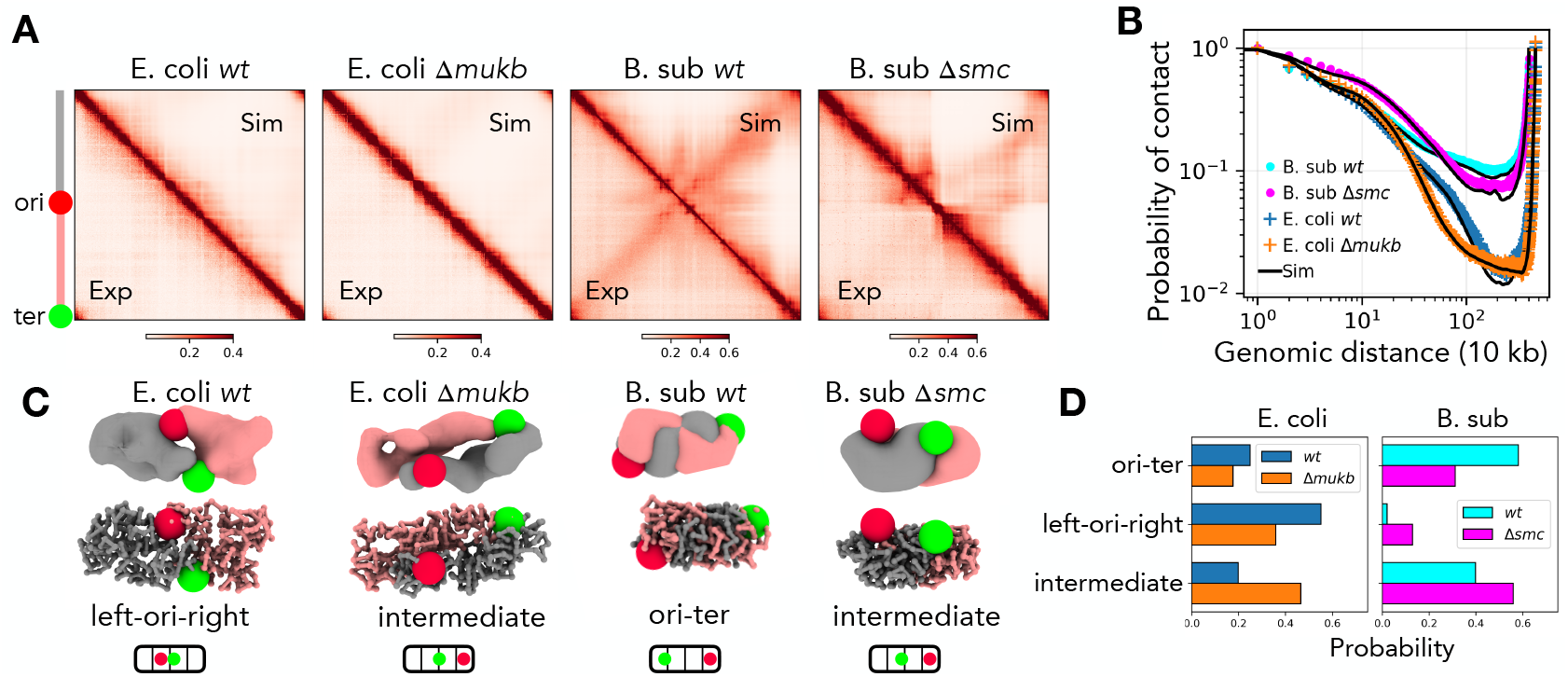
Optimized ensembles before replication. (A) Simulated (upper) and experimental (lower) Hi-C maps for `wt` and SMC-deficient mutants *E. coli* Δ`mukb` and *B. subtilis* Δ`smc` . The ori is located at the middle and ter at the end. (B) Mean Hi-C contact probability as a function of genomic distance, showing SMC-dependent enhancement of long-range ∼ 1 Mb, and suppression of short-range contacts ∼ 0.1 Mb. (C) Representative 3D structures showing ori in red and ter in green spheres. They illustrate three recurrent axial arrangements–left-ori-right, ori-ter, and intermediate states. The ori-ter configuration must have a distance between ori and ter that is greater than half the nucleoid length and at least one of the loci occupying a polar quarter. The left-ori-right organization must have their ori and ter residing in the mid quarters with an inter-locus distance less than a nucleoid quarter. All other configurations are intermediates. Each structure is shown in two representations, density-based surface construction (top) and polymer trace (bottom). Chromosome arms are shown in gray and salmon. (D) Probability of each axial organization in the learned structural ensembles, showing SMC loss perturbs their canonical `wt` configurations, increasing the proportion of intermediate states.

**FIG. 3.**
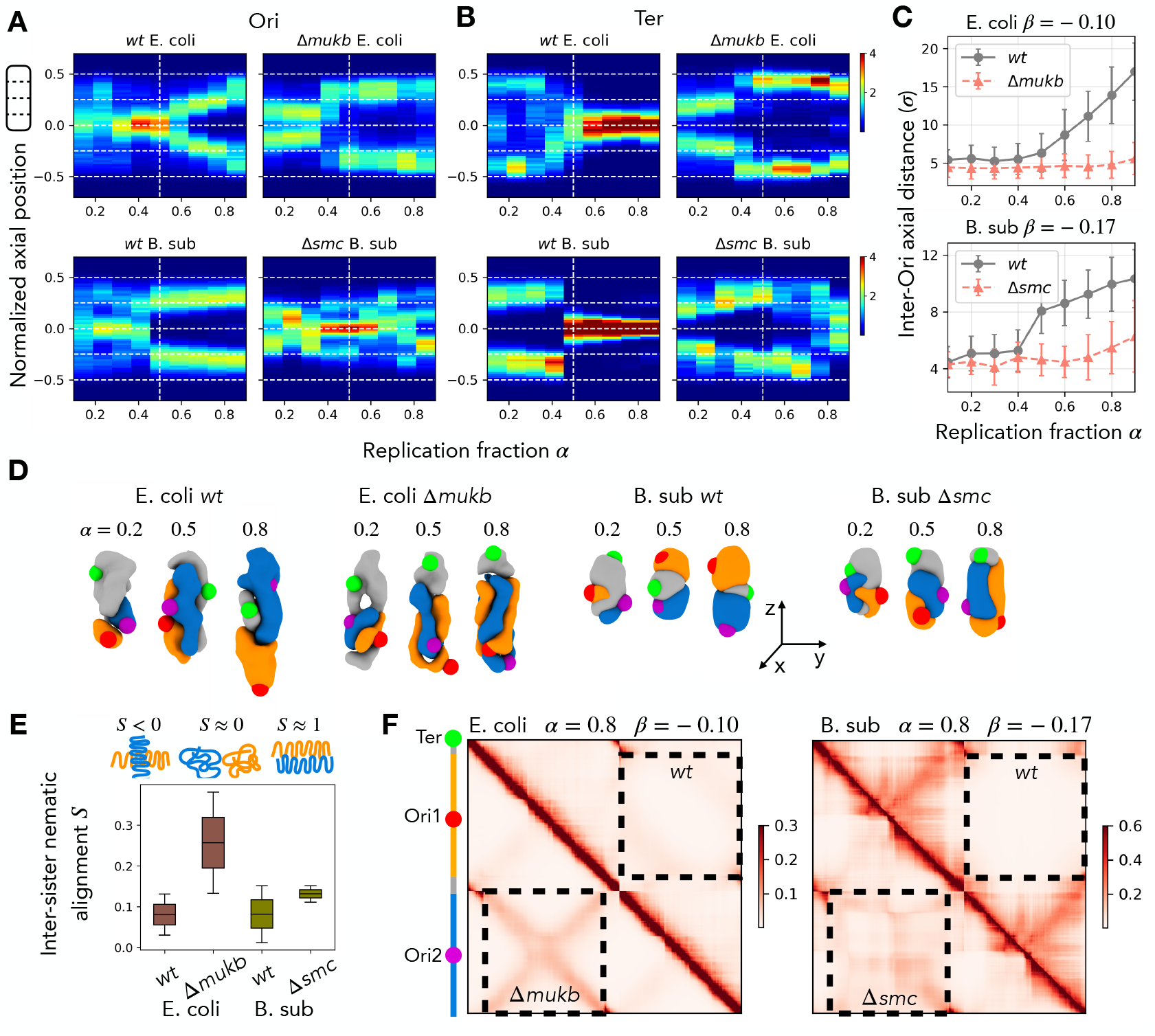
Structural transitions, segregation dynamics, and inter-sister interactions during replication. Kymographs showing the cell cycle axial dynamics of (A) the two ori and (B) ter across replication fractions *α* for *E. coli* (top) and *B. subtilis* (bottom). The corresponding mutants SMC-deficient *E. coli* and *B. subtilis* are shown in adjacent subpanels. The white horizontal lines denote the nucleoid quarters and the vertical white line denotes mid-replication. `wt` chromosomes undergo a mid-replication transition in which the ter moves toward the nucleoid center and the two ori separate toward opposite poles, whereas mutants lacking SMCs fail to undergo this rearrangement. (C) Inter-ori axial distance as a function of replication fraction, showing ori segregation in `wt` but proximal ori pair in Δ`mukb` and Δ`smc` mutants. (D) Representative 3D structures at early (*α* = 0.2), mid (*α* = 0.5), and late (*α* = 0.8) replication, illustrating `wt` ori splitting and ter repositioning versus persistent overlap and nematic alignment in mutants. Two ori and ter are red, purple, and green spheres, respectively. Chromosomes shown in density representation with two sisters in orange and blue, and unreplicated DNA in gray. (E) Inter-sister nematic alignment parameter *S* for *α* = 0.8, where *S* varies from −1*/*2 for perpendicular, to 0 for random isotropic, to 1 for perfectly aligned polymer. `wt` demonstrates more isotropic organization compared to SMC mutants with strong nematic alignment (*S >* 0.2) of replicated arms. (F) Simulated sister-chromosome–resolved Hi-C maps at *α* = 0.8 for *E. coli* and *B. subtilis*. The upper triangle shows `wt` versus SMC mutants Δ`mukb` and Δ`smc` in the lower triangles; inter-sister interaction blocks are highlighted by dashed rectangular boxes. SMC mutants show an increase in overall inter-sister contacts impeding segregation. The unsegregated state has nematic alignment that generates an X-shape in the inter-sister contact map. Together, these analyses show that SMC-mediated long-range compaction drives the structural transitions necessary for robust co-replication segregation.

**FIG. 4.**
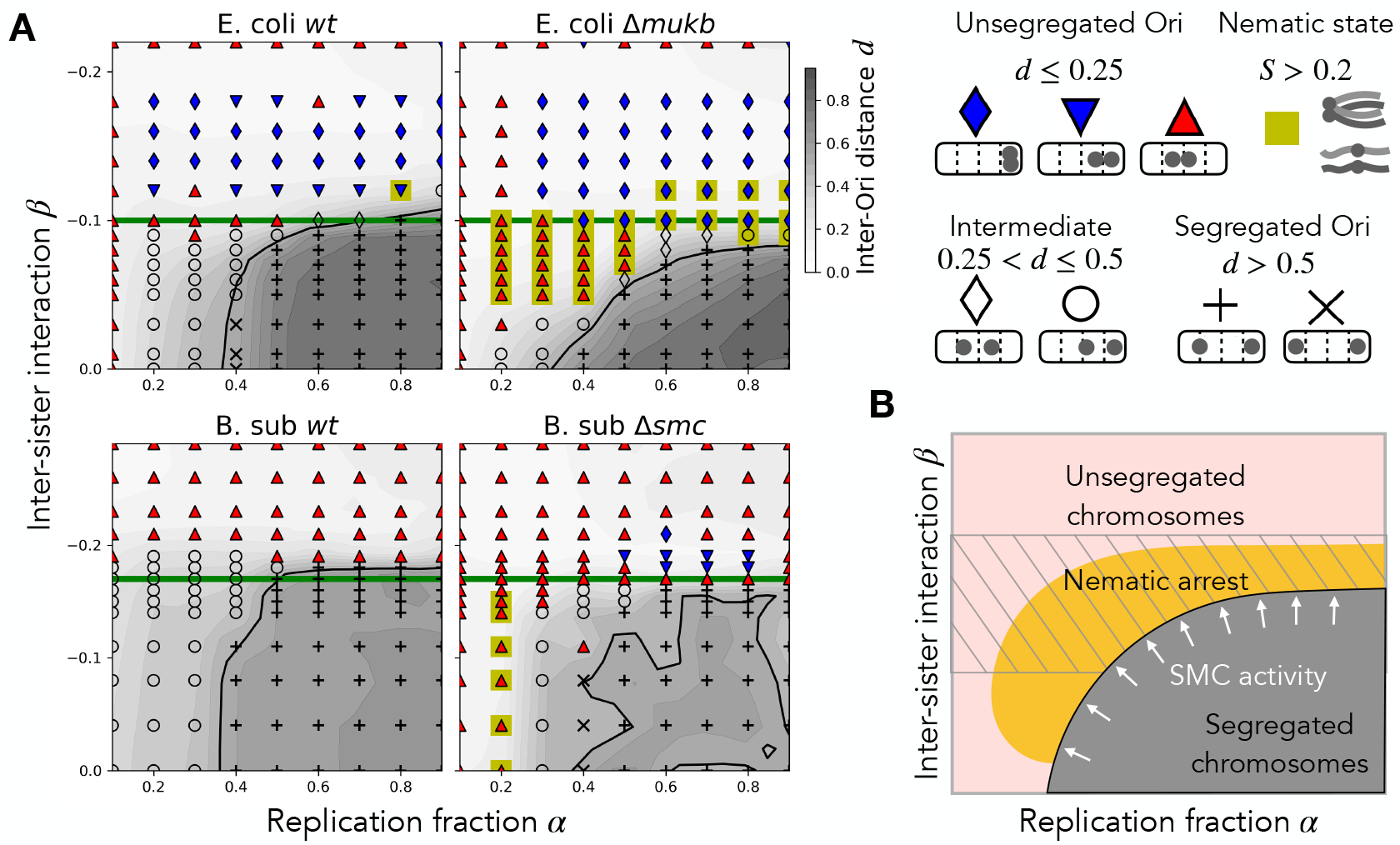
Regimes of chromosome segregation in rod-shaped bacteria revealed by energy-landscape modeling. (A) Segregation outcomes for *E. coli* (top) and *B. subtilis* (bottom) as a function of replication fraction *α* (horizontal axis) and mean inter-sister adhesion *β* (vertical axis). For each parameter pair, structural ensembles were generated under the learned chromosome landscapes for `wt` and SMC mutants (Δ`mukb` in *E. coli* and Δ`smc` in *B. subtilis*). Symbols denote the resulting segregation state of the two replicated origins–unsegregated (blue and red triangles, and blue diamonds), Intermediate (unfilled circles and diamonds), and segregated (gray crosses). Segregated means the inter-ori axial distance is larger than half the nucleoid length (*d >* 0.5). The background shading shows *d*, the mean inter-ori axial distance normalized to total nucleoid length, where darker regions correspond to larger separations. The black solid contour line represents inter-ori distance equal to half the nucleoid length *d* = 0.5; the thick horizontal green line corresponds to the non-specific baselines interaction, see Fig. 3. The yellow squares represent nematically aligned configurations in which sister chromosome arms lie parallel, a signature of unsegregated chromosomes. `wt` cells segregate their origins robustly over a wide range of adhesion strengths and replication stages, whereas Δ`mukb` and Δ`smc` mutants lose segregation competence at moderate *β* and instead enter a nematic-alignment regime. This nematic arrest emerges despite the inter-sister potential being structureless, reflecting a geometric consequence of the altered lengthwise compaction in SMC-deficient chromosomes. At higher adhesion values, both WT and mutant chromosomes collapse, leading to complete segregation failure. (B) Schematic phase diagram summarizing the three physically distinct regimes uncovered in panel A, where the cross-hatched region is physiologically relevant range of interactions learned from Hi-C. As replication fraction increases, the increased entropy of the sister chromosomes will favor segregation. However, biological activity introduces non-specific adhesive interactions *β* that reduce the entropic pressure to segregate. By establishing long-range intra-sister contacts, SMC activity enhances the effective segregation pressure. SMC mutants show loss of long-range compaction leading to significant overlap and emergence of nematic order that impedes segregation. At high adhesion, nonspecific inter-sister attractions dominate polymeric forces, causing chromosomes to collapse regardless of SMC activity. The diagram illustrates how SMC activity shifts the segregation boundary upward in *β* and enlarges the region of successful chromosome partitioning. Together, these results establish SMC complexes as key physical regulators of co-replication segregation, expanding the conditions under which rod-shaped bacteria reliably resolve their replicated genomes.

## I. MODEL

We develop a coarse-grained model of bacterial chromosomes in which each monomer represents 10 kb of DNA and evolves under Langevin dynamics within an effective energy landscape (Fig. 1 A) (see Eqs. (A1) and (A2) in Appendix). The polymer landscape corresponds to a ring homopolymer that includes nearest-neighbor harmonic bonds ((A3)) and soft excluded-volume interactions ((A4)). These terms alone are insufficient to reproduce experimentally observed chromosome architecture; hence, following previous studies [9–11], we introduce an additional pairwise bias potential 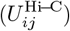 to be calibrated using Hi-C data ((B1)).

To infer 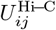, we employ an iterative maximum-entropy scheme in which simulated Hi-C contact probabilities ⟨*ϕ*⟩ are compared to experimentally measured probabilities to compute the loss gradient and update the bias potential. To convert distances into contact probabilities, we use a sigmoid *ϕ*(*r*) that is parameterized combining Hi-C and imaging studies (Fig. 1 B, (B2)). During optimization, the mean error between simulated and experimental Hi-C maps decreases monotonically and converges after ∼ 15 epochs (Fig. 1 C), at which point we take the learned 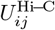 as the optimized landscape. This procedure is performed for *B. subtilis* and *E. coli* wild-type (`wt`) chromosomes and mutants, such as Δ`smc` and Δ`mukb` . Because the experimental Hi-C maps used in this optimization are obtained from asynchronous, exponentially growing populations and therefore average over cells at different stages of replication, we verified that this averaging does not significantly affect the inferred interaction landscapes.

The optimized interaction landscape reproduces Hi-C contact probabilities but yield globular conformations lacking the experimentally observed cylindrical shape. We therefore introduce a cylindrical confinement ((C1)) that constrains only the cross-section to experimentally measured nucleoid dimensions for *E. coli* and *B. subtilis* [15, 16]. This confinement minimally affects the Hi-C map while correctly reproducing axial dimensions and is kept fixed for all downstream modeling

The effective single-chromosome landscapes are then used to predict the architecture of partially replicated sister chromosomes (Fig. 1A). Each replication stage is parameterized by the replication fraction *α*, defined as the ratio of replicated DNA to the full chromosome length *L*; *α* = 0.5 corresponds to mid-replication. For a given *α*, we create a replicated sister-chromosome segment of length *αL* with the two sister origins positioned at its center. The two ends of this replicated segment, the replication forks, are harmonically bonded to the un-replicated circular chromosome, ensuring symmetric connectivity. Excluded-volume and confinement terms are identical for all monomers.

Because Hi-C cannot distinguish between the two nearly identical sister chromosomes, the intra-sister and sister-to-unreplicated interactions learned for the single chromosome were assumed to be symmetric. All components of the landscape can therefore be inherited directly from the single-chromosome model, except for the inter-sister component of the Hi-C bias potential, denoted 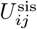 (Fig. 1 A). Importantly, using the learned intra-sister interactions for 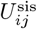 would incorrectly embed polymer-connectivity effects such as loop extrusion into interactions between different chromosomal copies. Instead, we interpret 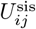 as representing adhesive forces arising from protein-mediated crosslinking, crowding, and other non-specific interactions in the dense nucleoid.

To model such nonspecific adhesion without imposing structure, we draw 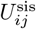 from a normal distribution with mean 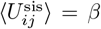 and a standard deviation 0.2|*β*| . The physiologically relevant estimate of *β* can be made from the baseline nonspecific interactions that generate low-probability long-range contacts in the learned single-chromosome landscape, such as the ori with ter interactions. This yields *β* = − 0.17 for *B. subtilis* and *β* = − 0.10 for *E. coli*, reflecting a stronger nonspecific compaction in *B. subtilis*. We first analyze sister-chromosome segregation using these inferred baseline interaction strengths (Fig. 3), and subsequently vary *β* to map the physiological regimes of chromosome segregation across species (Fig. 4).

## II. RESULTS

### A. Hi-C-derived structural ensembles validate imaging observations

We first confirm that the learned energy landscapes reproduce Hi-C contact patterns and known axial organizations, establishing a validated base model for all downstream analyses (Fig. 2 A). Both the SMC-deficient Δ`mukb` and Δ`smc` ensembles exhibit depletion of long-range contacts ∼0.5 − 1 Mb relative to `wt` and an enhancement in the short-range (Fig. 2 B). While SMC-mediated loop extrusion naturally explains the enhanced long-range contacts in `wt`, the mechanism increasing short-range compaction likely relates to NAP redistribution and altered transcription upon SMC depletion. Nevertheless, our inferred structural ensembles recapitulate these characteristic features.

A notable outcome of the learned landscapes is that constraining pairwise interactions and cylindrical cross-section is sufficient to break the symmetry of axial positioning (Fig. 2 C). As expected, our ensemble predictions preserve the reflection symmetry across the XY plane. We divide the nucleoid axial length into two *mid* and two *polar* quarters, and classify conformations as *ori-ter* and *left-ori-right* based on the positioning of ori and ter in these quarters. The ori-ter conformation is defined by an ori-to-ter distance greater than half the nucleoid length and at least one locus in a polar quarter. The left-ori-right conformations have ori and ter residing in the mid quarters with their inter-locus distance less than a nucleoid quarter. All other conformations are *intermediates*.

The *E. coli* `wt` ensemble shows the left–ori–right organization; whereas Δ`mukb` exhibits intermediate states with asymmetric ori-ter patterns (Fig. 2 C-D), as has been observed in imaging experiments [7, 30]. The *B. subtilis* `wt` landscape prefers the ori–ter pattern, while the mutant Δ`smc` shows intermediate states. Experiments suggest that *B. subtilis* chromosomes may adopt either conformations [14]; the predicted configurations are majorly ori-ter with modest contributions from intermediate states where ori and ter occupy neighboring quarters of the nucleoid (Fig. 2 C-D).

### B. Wild-type origins segregate mid-replication, accompanied by repositioning of the terminus towards nucleoid center

We next examine how chromosome architecture evolves during replication and identify a robust mid-replication structural transition that drives ori segregation and ter repositioning (Fig. 3). Using the optimized single-chromosome landscapes, we constructed effective potentials and corresponding structural ensembles for sister chromosomes across replication fractions, fixing the mean inter-sister interaction *β* to their respective baselines for *E. coli* and *B. subtilis*. We define the inter-ori axial distance normalized to the nucleoid length *d*, and use *d >* 0.5 as a criterion for ori segregation.

In the `wt` of both *E. coli* and *B. subtilis*, early replication stages (*α <* 0.3) show the two unsplit ori fluctuating within the two mid quarters, while the ter resides in a polar quarter (Fig. 3 A). The entry to the mid-replication stage (*α* ≈ 0.5) is marked by a stable mid-quarter localization of both ori, a conformation with elevated segregation pressure due to the increased overlap between sister chromosomes. During mid-replication, the ter and the adjoining ori swap positions, producing an ori–ter–ori arrangement with low inter-sister overlap and a marked increase in inter-ori separation (Fig. 3 B). This transition, essential for positioning the sister chromosomes at opposite cell halves during late replication (*α >* 0.7),has been experimentally observed in `wt` cells (Fig. 3 C) [15, 16].

In contrast, the SMC mutants Δ`mukb` and Δ`smc` lack the characteristic mid-replication swap; instead, the ter remains polar while the two unsplit ori either fluctuate within the mid quarters (Δ`smc`) or occupy the same pole (Δ`mukb`). This leads to unsegregated sister chromosomes and unresolved origins in late replication (Fig. 3 A-C). In `wt`, the enhancement of long-range intra-sister contacts effectively introduces an entropically driven inter-sister repulsion that underlies the mid-replication transition; in the mutants, this repulsion is weakened, impairing segregation.

### C. Unsegregated sister chromosomes of SMC mutants exhibit emergent nematic order with distinctive Hi-C signatures

We then analyze the SMC-deficient mutants and show that the loss of long-range compaction produces nematic alignment of sister chromosomes, which resists segregation and predicts a testable inter-sister Hi-C signature (Fig. 3 F). We use the term nematic alignment in a descriptive sense to denote strong parallel orientation of sister chromosome arms. We quantify this ordering using the nematic-like order parameter *S* ((D1)), that ranges from −1/2 for perpendicular configurations to 1 for perfectly aligned arms ((D1)). Sister chromosomes coarse-grained by 30 monomers when show *S >* 0.2 are classified to have a significant nematic alignment (Figs. 3 E).

In extrusion-deficient mutants, redistribution of lengthwise compaction causes the two sister-chromosome arms to behave as two U-shaped rods with orthogonally oriented planes that exhibit higher nematic order (Fig. 3 D-E). This low-entropy configuration resists segregation of sister chromosomes, and in some cases, results in a novel X-shaped pattern in the inter-sister block of the simulated Hi-C maps (Fig. 3 E). The *B. subtilis* chromosomes exhibited lower nematic alignment than *E. coli* due to their higher overall compaction. Importantly, the inter-sister interaction potential itself is structureless, its elements are randomly drawn around the mean *β*, so the observed nematic alignment and Hi-C signature is an emergent geometric consequence of altered lengthwise compaction rather than an imposed interaction pattern. Traditionally, Hi-C does not resolve sister chromosomes and is incapable of detecting signatures of inter-sister organization [12, 13]; however, improved Hi-C technology, focused on distinguishing sister-chromosomes [31, 32], should be able to verify the model predictions.

### D. Structural organization by SMC proteins enhances robustness of chromosome segregation

To test how SMC-dependent architecture modulates segregation fidelity, we varied the mean inter-sister adhesion *β* and mapped the regimes in which `wt` and mutant chromosomes successfully segregate. Our earlier results showed that SMC-mediated structural organization can drive co-replication segregation when *β* is comparable to a typical nonspecific interaction. Physically, *β* represents the combined effects of nonspecific nucleoid crosslinkers, homology-mediated pairing, NAP-driven protein bridging, and crowding-enhanced attractions, and may vary during the cell cycle in response to environmental or regulatory cues. Importantly, we posit, SMCs do not contribute to *β*, because of their strictly intra-chromosome-specific activity. We therefore varied *β* across a wide range to test the robustness of SMC-driven segregation across different replication fractions *α* (Fig. 4).

We scanned *β* from zero, corresponding to an absence of inter-sister adhesion, to values representing strongly attractive interactions in the learned single-chromosome potentials. Because `wt` *E. coli* and *B. subtilis* exhibit different baseline interaction strengths, the relevant *β* ranges differ across species (Fig. 4 A, solid green line). Despite this, both organisms display similar segregation regimes when comparing `wt` and SMC mutants (Fig. 4 B), indicating a conserved physical mechanism.

When inter-sister adhesion is weak, the polymer entropy of the long replicated arms is sufficient to separate the sisters during late replication, even in SMC-deficient mutants. In this regime of high *α* and low *β*, both the `wt` and mutant ensembles show inter-ori distances larger than half the nucleoid length (*d >* 0.5), establishing segregated ori. As *β* increases to moderate values, `wt` chromosomes continue to undergo the characteristic mid-replication transition and segregate their ori, whereas SMC mutants fail to do so and instead develop nematic alignment of unresolved sister arms. At high *β*, adhesion dominates the ensembles and both `wt` and mutants fail to segregate their origins. These results show that SMC-mediated long-range lengthwise compaction broadens the range of inter-sister adhesion strengths over which ori segregates faithfully, buffering chromosome segregation against variable adhesive forces across diverse cellular contexts (Fig. 4 B).

### E. Nucleoid-associated proteins interact with SMCs to regulate ori segregation

Finally, we investigate how key nucleoid-associated proteins (NAPs) modulate SMC-dependent chromosome organization and find that they tune, but do not replace, the core SMC-driven segregation mechanism. We examined two well-studied NAPs—MatP in *E. coli* [13] and ParB in *B. subtilis* [12]—to assess their impact on chromosome architecture and sister-chromosome segregation.

MatP in *E. coli* binds the terminus-proximal region and excludes the extrusion complex MukBEF from the ter domain. Consistent with this, MatP mutants (Δ`matp`) exhibit a ter-specific increase in long-range contacts (0.5 Mb), likely reflecting elevated MukBEF activity near the terminus [13, 33]. The optimized Δ`matp` models adopt an ori–ter arrangement and predict enhanced segregation of ter-proximal regions, suggesting that MatP normally suppresses premature ter segregation. In contrast, deleting both MukB and MatP (Δ`mukbmatp`) shows a severe loss of segregation robustness, including in the ter region, indicating that the increased ter segregation observed in Δ`matp` arises from residual MukBEF activity.

ParB is part of the ParABS partitioning system in *B. subtilis* that loads SMCs onto origin-proximal DNA, driving the Hi-C counter-diagonal observed in `wt` cells [12, 16, 24, 34, 35]. Consistently, disruption of ParB eliminates the counter-diagonal while preserving long-range contacts [12, 16, 34]. To test whether ParB is required for sister-chromosome segregation, we optimized the energy landscape of ParB-deletion mutants (Δ`par`) [12]. The resulting single-chromosome structures predominantly exhibited a left–ori–right configuration. The Δ`par` mutant displayed larger inter-ori distances during early–mid replication than `wt`, yet ultimately segregated their sister chromosomes less efficiently in late replication. Additionally, the Δ`par` ensembles showed no nematic alignment of the sister chromosome, reiterating the loss of SMC-mediated long-range lengthwise compaction as the major driver of nematic order.

Together, these results indicate that while NAPs such as MatP and ParB interact with SMCs to reshape local chromosomal organization, particularly around ori and ter, the fundamental structural transitions that drive ori segregation are primarily mediated by SMC complexes, even in the absence of their localized loading.

## III. DISCUSSION

We present an energy-landscape framework that provides a quantitative, mechanistic account of how SMC complexes shape bacterial chromosome organization to ensure faithful co-replication segregation. By integrating Hi-C-derived constraints with coarse-grained polymer physics, we show that SMC lengthwise-compaction activity reorganizes the nucleoid during replication by enhancing long-range and suppressing short-range contacts. This SMC-driven remodeling produces a cooperative mid-replication transition in which the terminus moves towards the center while replicated origins separate toward opposite cell poles, offering a unifying physical explanation for the spatiotemporal choreography of bacterial chromosomes across species.

Our approach constructs a least-biased effective-equilibrium ensemble that reproduces the measured Hi-C contact landscape while remaining agnostic to the microscopic processes that generate these contacts (Figs. 1 and 2). Active mechanisms such as loop extrusion enter implicitly through their influence on observed contact probabilities, rather than through explicit dynamical modeling [17–22, 24–26]. Recent polymer-physics models have shown that loop extrusion can direct entropic forces to promote bacterial chromosome segregation, while passive loop formation and confinement can also facilitate segregation by modifying polymer overlap and entropy [24, 36].

Replication is treated as a sequence of discrete stages, essentially assuming that the primary effect of the replication apparatus is to perturb short-range fluctuations near the fork while large-scale architecture is determined chiefly by DNA length and global structural constraints. Recent studies modeling bacterial co-replication segregation [24–26] have considered both a steady-state approach like ours and a dynamic approach with explicit replication-progression events, and shown that both yield the same segregation dynamics. Polymer-based models have further shown that DNA replication itself can serve as an active driver of large-scale genome reorganization, independent of SMC-mediated loop extrusion. [37–39]. In these dynamic descriptions, replication forks can act as barriers to loop extrusion and introduce fork-associated interactions; in our framework, such replication-specific effects are absorbed into the effective interaction landscape inferred from Hi-C. This coarse-grained description allows us to focus on robust, large-scale architectural constraints and to track how these constraints shift across genetic backgrounds, isolating the dominant physical mechanisms shaping chromosome organization.

Additional constraints, such as tethering of genomic segments (ori or ter) to the cell poles by specific NAPs may arise during the cell cycle [7, 12]. Our model effectively encodes them via the pairwise contacts; substantial disagreements would suggest the need to explicitly incorporate such constraints. However, as far as axial localizations during exponential growth in *E. coli* and *B. subtilis* go, our cylindrically confined pairwise-interaction model quantitatively captures the observed phenomenology [14–16].

The polymer structural ensembles, characterized by soft self-exclusion between all monomers, allow polymer-topology fluctuations. This accounts for the double-strand-passing activity of Type-II DNA topoisomerase enzymes, such as bacterial gyrase and Topo IV [7]. In such fluctuating topology ensembles, contact enrichment embeds topological entanglement [5]. This suggests that the enriched contacts between the sister chromosome arms in early-to-mid replication, preceding ori splitting, are likely to have higher entanglements and require DNA topoisomerase activity to execute the mid-replication transition. Explicit dynamic modeling of strand passage by topoisomerase enzymes has observed their importance in preventing catastrophic DNA entanglements during both bacterial [25, 26] and eukaryotic replication [37]. Experiments on bacterial systems suggest an interplay of SMC complexes and Topoisomerase enzymes in regulating sister-chromosome segregation dynamics [40, 41].

Comparison of `wt` and SMC-deficient mutants Δ`mukb` and Δ`smc` highlights how the redistribution of length-wise compaction acts as the principal physical force co-ordinating chromosome partitioning. Loss of long-range compaction in SMC mutants results in nematic-like alignment of sister chromosomes that restricts origin segregation. The nematic state may be observed in sister-chromosome resolved HiC [31, 32] via specific inter-sister Hi-C signatures (Fig. 3). The nematic alignment is further reinforced by enhanced short-range compaction in SMC mutants (Fig. 2 B), which likely reflects subtleties of SMC function. In light of recent works suggesting that transcription-induced loops or supercoiled domains may drive local compaction state [42], the short-range compaction may be a result of altered transcription and NAP distribution in the absence of SMCs. Further modeling, incorporating aspects of DNA mechanics [25, 26] and transcription dynamics [43], will be required to disentangle SMC loop extrusion from transcription-mediated genome restructuring.

By systematically varying the effective inter-sister adhesion, we show that SMC activity broadens the parameter regime in which segregation is robust, buffering against adhesive interactions in the crowded bacterial nucleoid (Fig. 4). Importantly, such effective adhesion likely arises from multiple, non-exclusive sources, including protein-mediated crosslinking and nonspecific crowding effects that may act both within and between sister chromosomes. This robustness supports the view that SMC-mediated organization provides an evolutionarily conserved strategy for promoting chromosome segregation in an adhesive intracellular environment. Future experimental approaches that combine spatial imaging with contact-based measurements may help further constrain the dominant physical contributors to inter-sister interactions.

Finally, our findings provide a framework to understand how NAPs interact with SMCs to regulate chromosome architecture. While ParB- or MatP-dependent effects reshape local structures, such as ori-proximal extrusion in *B. subtilis* or MukBEF exclusion from ter in *E. coli*, the core SMC-driven mid-replication transition persists in their absence. Thus, NAPs tune but do not replace the fundamental SMC-dependent mechanism of segregation via lengthwise compaction. This suggests that certain nucleoid-associated proteins, such as MatP, may be dispensable for robust large-scale chromosome segregation under the conditions studied, an insight with potential implications for synthetic biology efforts aimed at constructing minimal cellular systems [44]. Together, our data-driven models generate testable predictions for imaging and Hi-C experiments, establishing a generalizable physical paradigm for chromosome organization and segregation across bacteria.

## Appendix A: Simulating chromosome dynamics using polymer potentials

We integrate the following Langevin equation of motion to get the dynamics of each coarse-grained monomers.

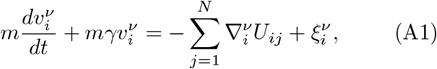

where the mass of each monomer is *m* = 1.0 and the friction coefficient is *γ* = 0.1. The velocity components are 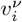, where ν ∈ (*x, y, z*) correspond to three cartesian coordinates and *i, j* ∈ (1,, *N*) for particles. The thermal noise, given by 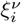, has zero mean and is uncorrelated: 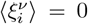 and 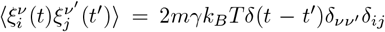, where *k*_*B*_*T* = 1.0 sets the scale for thermal energy with *T* denoting the temperature and *k*_*B*_ the Boltzmann constant. The velocity components are integrated via the leap-frog method with a time step *dt* = 0.01. The term 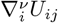 refers to the forces acting along ν on the *i*-th particle, derived from the pairwise chromosome potential *U*_*ij*_, written as follows.

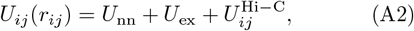

where 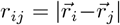, is the inter-particle distance, and the terms on the R.H.S. are the contributions from, respectively, the nearest neighbor connections of the polymer, soft self-exclusion, and the data-driven pairwise interactions learned from Hi-C.

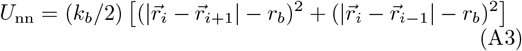

The nearest neighbor interactions are harmonic springs, *U*_nn_ with an equilibrium distance *r*_*b*_ = 1.0*σ*, where *σ*, the monomer diameter, is the unit of distance, and bond stiffness *k*_*b*_ = 30.0*k*_*B*_*T/σ*^2^. The same potential is used for circularization of the chromosome polymer by connecting the last and the first monomers. The self-avoidance potential *U*_ex_ is a soft-repulsive sigmoid,

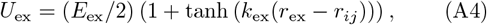

where the slope of the sigmoid is controlled by *k*_ex_ = 5.0*/σ*, the cutoff distance *r*_ex_ = 0.7*σ*, and the maximum strength of the self-avoidance is given by *E*_ex_ = 4.0*k*_*B*_*T* . Finally, the potential *U* ^Hi−C^ is derived from the pairwise constraints of the Hi-C data, as described in the next section. We use all the potentials defined above as the reference on top of which *U* ^Hi−C^ is learned to recapitulate the experimental Hi-C maps in simulation ensembles.

We initialized the polymer using a random gas-like conformation and then equilibrated under the chromosome potential (Eq. A2) for 10^6^ simulation steps before storing data for calculations. We ran 20 independent replicates for each parameter sets, with each replicate run spanning 5 × 10^6^ steps. The simulations used a time step of *dt* = 0.01 and the structures were sampled every 500 steps, giving a total of 2 × 10^5^ structures per replicate run. All the structures from the replicates were combined to do the statistical analyses and computation of simulated Hi-C maps.

Chromosome simulations were performed using the freely available custom software *chromdyn*, inspired by OpenMiChroM [45] and implemented as a wrapper around OpenMM [46]. We used VMD with Trace, VDW, and QuickSurf representations to generate the polymer visualizations [47].

## Appendix B: Optimizing the chromosome potential using Hi-C data

We seek to find the potential 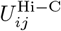, such that a polymer equilibrated under the total potential *U* ((A2)) recapitulates the experimental Hi-C maps [12, 13]. The problem may be cast as a search for a probability distribution such that samples from the distribution give chromosome structures consistent with the Hi-C maps. We build upon a statistical mechanical approach of entropy maximization, originally proposed by E.T. Jaynes [8] and later implemented in various forms for inferring chromosome structures [9–11].

The most general potential consistent with the maximum-entropy approach gives

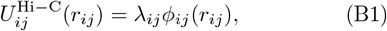

The coefficient *λ*_*ij*_ is the Lagrange multiplier associated with the Hi-C contact probability between the monomer pairs *i* and *j*, denoted by *ϕ*_*ij*_ ((B2)). Note we consider only the interactions between non-neighboring loci pairs, i.e., *j > i*+1, since the nearest neighbor contacts are fixed by bonded interactions ((A3)). Using the Lagrangian as the loss function, 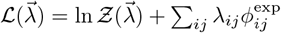,where *Ƶ* is the partition function, the error gradient takes a simple analytical form, 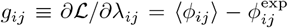. We use the gradient-descent update, *λ*(*t* + 1) = *λ*(*t*) + *ηg*_*ij*_(*t*), with a fixed learning rate *η* = 0.2, where *t* represents the training epoch.

The experimental Hi-C maps used for inference were obtained from asynchronous, exponentially growing populations and therefore represent averages over multiple replication states. We explicitly tested the robustness of the inferred interaction potentials to this replication-state averaging. Incorporating replication-stage averaging had only a minimal effect on the optimized chromosome potentials, resulting in deviations of approximately 3% relative to those learned from non-replicating chromosomes.

### Preprocessing experimental Hi-C maps for optimization

We normalized the experimental Hi-C maps by dividing all the matrix elements by the mean nearest-neighbor values, which maximized the experimental probability of contact with the nearest neighbor, *P*_*i,i*+1_ = 1.0. This condition is always satisfied in the simulated ensembles because of the bonded neighbors. While the provided *B. subtilis* Hi-C matrices were of 10 kb resolution [12], the *E. coli* Hi-C were of 5 kb resolution [13]. We used block-averaging downsampling by a factor of 2 to obtain 10 kb *E. coli* Hi-C maps.

### Converting 3D distances to Hi-C probability

We define a Hi-C contact from 3D distance between loci pairs *i* and *j* using a modified sigmoid as follows.

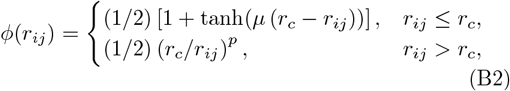

where, we used *µ* = 2.0*/σ, r*_*c*_ = 2.0*σ*, and *p* = 4.0, which showed good agreement with imaging data (Fig. 1 B). Note that the power-law tail of the function, that relaxes conformational constraints for low-probability contacts between far-away particles *r*_*ij*_ *> r*_*c*_, is only used for the simulated Hi-C computation. The contacts driving the potentials do not use this long-range decay, making all the potentials purely physical contact-based.

## Appendix C: Predicting structural ensembles of sister chromosomes

To create the geometry of sister chromosomes at replication fraction *α*, we add a sister chromosome segment of length *αL* to a circular chromosome of size *L* by harmonically bonding the two extremities of the sister chromosome to the two monomers representing the replication fork positions on the two chromosome arms. Except for the component of the inter-sister interactions 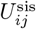, all the other potentials are directly derived from the learned single chromosome potential using symmetry (Fig. 1 A). We hypothesize the inter-sister interaction to be non-specific, hence draw elements of 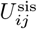 from a random distribution. We vary the non-specific interaction strength by varying the mean of the random distribution 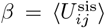. The width of the distribution is set at 0.2|*β*| to add moderate noise.

We additionally implement a cylindrically symmetric confinement potential that kicks in when the cross-sectional radius is larger than *ρ*_conf_.

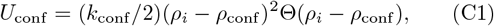

where, 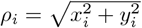 is the perpendicular distance from the *z*-axis, *k*_conf_ = 10.0*k*_*B*_*T/σ*^2^ is the stiffness of the confinement potential, and the Heavyside-theta function with the usual definition: 1 if *ρ*_*i*_ *> ρ*_conf_, else 0. The confinement-free optimized potential produces globular, rather than cylindrical chromosomes, as observed in bacteria [15, 16]. Adding this radial confinement bias on top of the learned potentials reproduced the observed aspect ratio of bacterial nucleoids. We used *ρ*_conf_ = 4.0*σ* and 2.5*σ* for *E. coli* and *B. subtilis*, respectively.

## Appendix D: Nematic alignment of sister chromosomes

We defined the nematic order parameter

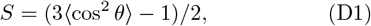

where *θ* is the angle between coarse-grained backbone tangent vectors, one from each sister chromosome, evaluated at loci that are equidistant along the chromosome contour from the replication fork. *S* = −1/2 for perpendicular configurations, *S* = 0 for random isotropic configurations, and *S* = 1 for perfectly aligned arms. The nematic alignment *S* increased monotonically and then saturated when the tangent vectors were constructed from an increasingly coarse-grained polymer. In Figs. 3 E and 4 A, we coarse-grain the sister chromosomes by 30 monomers (∼ 0.3 Mb).

## Data availability

The custom software for chromosome simulations and optimization chromdyn is freely available: https://github.com/s-brahmachari/chromdyn.git. All the experimental data used in this study are previously published and available from the Gene Expression Omnibus (GEO) repositories: Hi-C data of *B. subtilis* (GSM1671399, GSM1671401, GSM1671414), Hi-C data of *E. coli* (GSM2870416, GSM2870418, GSM2870421, GSM2870420). The imaging data for *B. subtilis* are from Ref. [12], and for *E. coli* are from Refs. [13, 29].

## Acknowledgements

We thank the National Science Foundation (Grants No. PHY-2019745, No. PHY2210291, and No. PHY-2014141) and the Welch Foundation (Grant No. C-1792) for funding. We gratefully acknowledge the Center for Research Computing (CRC) at Rice University for providing computational resources for this work. We also thank Advanced Micro Devices for donating critical hardware and support resources from its HPC Fund that contributed to this work.

## References

[1] C. Hoencamp and B. Rowland, Genome control by smc complexes, Nat Rev Mol Cell Biol 24, 633–650 (2023).

[2] E. Kim, R. Barth, and C. Dekker, Looping the genome with smc complexes, Annual Review of Biochemistry 92, 15 (2023).

[3] S. Nolivos and D. Sherratt, The bacterial chromosome: architecture and action of bacterial smc and smc-like complexes, FEMS Microbiology Reviews 38, 380–392 (2014).

[4] T. Hirano, Condensin-based chromosome organization from bacteria to vertebrates, Cell 164 (2016).

[5] S. Brahmachari and J. F. Marko, Chromosome disentanglement driven via optimal compaction of loop-extruded brush structures, Proceedings of the National Academy of Sciences 116, 201906355 (2019).

[6] C. Hoencamp, O. Dudchenko, A. M. Elbatsh, S. Brahmachari, J. A. Raaijmakers, T. van Schaik, Ángela Sedeño Cacciatore, V. G. Contessoto, R. G. van Heesbeen, B. van den Broek, A. N. Mhaskar, H. Teunissen, B. G. S. Hilaire, D. Weisz, A. D. Omer, M. Pham, Z. Colaric, Z. Yang, S. S. Rao, N. Mitra, C. Lui, W. Yao, R. Khan, L. L. Moroz, A. Kohn, J. S. Leger, A. Mena, K. Holcroft, M. C. Gambetta, F. Lim, E. Farley, N. Stein, A. Haddad, D. Chauss, A. S. Mutlu, M. C. Wang, N. D. Young, E. Hildebrandt, H. H. Cheng, C. J. Knight, T. L. Burnham, K. A. Hovel, A. J. Beel, P. J. Mattei, R. D. Kornberg, W. C. Warren, G. Cary, J.L. Gómez-Skarmeta, V. Hinman, K. Lindblad-Toh, F. D. Palma, K. Maeshima, A. S. Multani, S. Pathak, L. Nel-Themaat, R. R. Behringer, P. Kaur, R. H. Medema, B. van Steensel, E. de Wit, J. N. Onuchic, M. D. Pierro, E. L. Aiden, and B. D. Rowland, 3d genomics across the tree of life reveals condensin ii as a determinant of architecture type, Science 372, 10.1126/SCIENCE.ABE2218 (2021).

[7] A. Badrinarayanan, T. B. Le, and M. T. Laub, Bacterial chromosome organization and segregation., Annu Rev Cell Dev Biol. 31, 171 (2015).

[8] E. T. Jaynes, Information theory and statistical mechanics, Phys. Rev. 106, 620 (1957).

[9] B. Zhang and P. G. Wolynes, Topology, structures, and energy landscapes of human chromosomes, Proceedings of the National Academy of Sciences of the United States of America 112, 6062 (2015).

[10] M. D. Pierro, B. Zhang, E. L. Aiden, P. G. Wolynes, and J. N. Onuchic, Transferable model for chromosome architecture, Proceedings of the National Academy of Sciences of the United States of America 113, 12168 (2016).

[11] A. B. O. Junior, M. F. Mello, R. J. Oliveira, E. Dodero-Rojas, S. Brahmachari, V. G. Contessoto, and J. N. Onuchic, A data-driven chromatin model reveals spatial and dynamic features of genome organization, Proceedings of the National Academy of Sciences 123, e2530583123 (2026), https://www.pnas.org/doi/pdf/10.1073/pnas.2530583123.

[12] X. Wang, T. B. Le, B. R. Lajoie, J. Dekker, M. Laub, and D. Z. Rudner, Condensin promotes the juxtaposition of dna flanking its loading site in bacillus subtilis., Genes Dev. 29, 1661 (2015).

[13] V. S. Lioy, A. Cournac, M. Marbouty, S. Duigou, J. Mozziconacci, O. Espéli, F. Boccard, and R. Koszul, Multiscale structuring of the e. coli chromosome by nucleoid-associated and condensin proteins, Cell 172, 771 (2018).

[14] X. Wang, P. M. Llopis, and D. Z. Rudner, Bacillus subtilis chromosome organization oscillates between two distinct patterns, Proceedings of the National Academy of Sciences of the United States of America 111, 12877 (2014).

[15] J. K. Fisher, A. Bourniquel, G. Witz, B. Weiner, M. Prentiss, and N. Kleckner, Four-dimensional imaging of e. coli nucleoid organization and dynamics in living cells, Cell 153, 882 (2013).

[16] M. Marbouty, A. L. Gall, D. I. Cattoni, A. Cournac, A. Koh, J. B. Fiche, J. Mozziconacci, H. Murray, R. Koszul, and M. Nollmann, Condensin- and replication-mediated bacterial chromosome folding and origin condensation revealed by hi-c and super-resolution imaging, Molecular Cell 59, 588 (2015).

[17] E. Alipour and J. F. Marko, Self-organization of domain structures by dna-loop-extruding enzymes., Nucleic Acids Res. 40, 11202 (2012).

[18] A. L. Sanborn, S. S. Rao, S. C. Huang, N. C. Durand, M. H. Huntley, A. I. Jewett, I. D. Bochkov, D. Chinnappan, A. Cutkosky, J. Li, K. P. Geeting, A. Gnirke, A. Melnikov, D. McKenna, E. K. Stamenova, E. S. Lander, and E. L. Aiden, Chromatin extrusion explains key features of loop and domain formation in wild-type and engineered genomes, Proceedings of the National Academy of Sciences of the United States of America 112, E6456 (2015).

[19] A. Goloborodko, M. V. Imakaev, J. F. Marko, and L. Mirny, Compaction and segregation of sister chromatids via active loop extrusion, eLife 5, e14864 (2016).

[20] G. Fudenberg, M. Imakaev, C. Lu, A. Goloborodko, and L. A. Mirny, Formation of chromosomal domains by loop extrusion, Cell Rep. 15, 2038 (2016).

[21] E. J. Banigan, A. A. van den Berg, H. B. Brandao, J. F. Marko, and L. A. Mirny, Chromosome organization by one-sided and two-sided loop extrusion, eLife 9, 10.7554/eLife.53558 (2020).

[22] H. Salari and D. Jost, Loop extrusion provides mechanical robustness to chromatin 10.1101/2025.09.16.676638 (2025).

[23] S. Jun and B. Mulder, Entropy-driven spatial organization of highly confined polymers: Lessons for the bacterial chromosome (2006).

[24] J. Harju, M. van Teeseling, and C. Broedersz, Loop-extruders alter bacterial chromosome topology to direct entropic forces for segregation., Nat Commun 15 (2024).

[25] B. R. Gilbert, Z. R. Thornburg, T. A. Brier, J. A. Stevens, F. Grünewald, J. E. Stone, S. J. Marrink, and Z. Luthey-Schulten, Dynamics of chromosome organization in a minimal bacterial cell, Frontiers in Cell and Developmental Biology 11, 10.3389/fcell.2023.1214962 (2023).

[26] Z. R. Thornburg, A. Maytin, J. Kwon, T. A. Brier, B. R. Gilbert, E. Fu, Y.-L. Gao, J. Quenneville, T. Wu, H. Li, T. Long, W. Pezeshkian, L. Sun, J. I. Glass, A. Mehta, T. Ha, and Z. Luthey-Schulten, Bringing the genetically minimal cell to life on a computer in 4d, bioRxiv 10.1101/2025.06.10.658899 (2025).

[27] A. Wasim, A. Gupta, and J. Mondal, A hi–c data-integrated model elucidates e. coli chromosome’s multiscale organization at various replication stages, Nucleic Acids Research 49, 3077 (2021), https://academic.oup.com/nar/article-pdf/49/6/3077/36885066/gkab094σupplementalϕile.pdf.

[28] J. J. B. Messelink, M. C. F. van Teeseling, J. Janssen, M. Thanbichler, and C. P. Broedersz, Learning the distribution of single-cell chromosome conformations in bacteria reveals emergent order across genomic scales., Nat Commun 12, 10.1038/s41467-021-22189-x (2021).

[29] O. Espeli, R. Mercier, and F. Boccard, Dna dynamics vary according to macrodomain topography in the e. coli chromosome, Molecular Microbiology 68, 1418 (2008).

[30] O. Danilova, R. Reyes-Lamothe, M. Pinskaya, D. Sherratt, and C. Possoz, Mukb colocalizes with the oric region and is required for organization of the two escherichia coli chromosome arms into separate cell halves, Molecular Microbiology 65, 1485 (2007).

[31] M. Mitter, C. Gasser, Z. Takacs, C. C. Langer, W. Tang, G. Jessberger, C. T. Beales, E. Neuner, S. L. Ameres, J. M. Peters, A. Goloborodko, R. Micura, and D. W. Gerlich, Conformation of sister chromatids in the replicated human genome, Nature 586, 139 (2020).

[32] M. E. Oomen, A. K. Hedger, J. K. Watts, and J. Dekker, Detecting chromatin interactions between and along sister chromatids with sisterc, Nature Methods 17, 1002 (2020).

[33] J. Mäkelä and D. J. Sherratt, Organization of the escherichia coli chromosome by a mukbef axial core, Molecular Cell 78, 250 (2020).

[34] X. Wang, H. B. Brandão, T. B. K. Le, M. T. Laub, and D. Z. Rudner, Bacillus subtilis smc complexes juxtapose chromosome arms as they travel from origin to terminus, Science 355, 524 (2017).

[35] A. Anchimiuk, V. S. Lioy, F. P. Bock, A. Minnen, F. Boccard, and S. Gruber, A low smc flux avoids collisions and facilitates chromosome organization in Bacillus subtilis, ELife 10, e65467 (2021).

[36] D. Mitra, S. Pande, and A. Chatterji, Polymer architecture orchestrates the segregation and spatial organization of replicating e. coli chromosomes in slow growth, Soft Matter 18, 5615 (2022).

[37] D. D’Asaro, M. M. Tortora, C. Vaillant, J. M. Arbona, and D. Jost, Dna replication and polymer chain duplication reshape the genome in space and time, Physical Review X 14, 10.1103/PhysRevX.14.041020 (2024).

[38] G. Forte, S. Buonomo, P. R. Cook, N. Gilbert, D. Marenduzzo, and E. Orlandini, Modeling the 3d spatiotemporal organization of chromatin replication, PRX Life 2, 033014 (2024).

[39] D. D’Asaro, J.-M. Arbona, V. Piveteau, A. Piazza, C. Vaillant, and D. Jost, Genome-wide modeling of dna replication in space and time confirms the emergence of replication specific patterns in vivo in eukaryotes, Genome Biol. 26, 10.1186/s13059-025-03872-4 (2025).

[40] X. Wang, R. Reyes-Lamothe, and D. J. Sherratt, Modulation of escherichia coli sister chromosome cohesion by topoisomerase iv, Genes and Development 22, 2426 (2008).

[41] E. Nicolas, A. L. Upton, S. Uphoff, O. Henry, A. Badrinarayanan, and D. Sherratt, The smc complex mukbef recruits topoisomerase iv to the origin of replication region in live escherichia coli, mBio 5, 10.1128/mBio.01001-13 (2014).

[42] A. Bignaud, C. Cockram, C. Borde, J. Groseille, E. Allemand, A. Thierry, M. Marbouty, J. Mozziconacci, O. Espéli, and R. Koszul, Transcription-induced domains form the elementary constraining building blocks of bacterial chromosomes, Nature Structural and Molecular Biology 31, 489 (2024).

[43] S. Tripathi, S. Brahmachari, J. N. Onuchic, and H. Levine, DNA supercoiling-mediated collective behavior of co-transcribing RNA polymerases, Nucleic Acids Research 50, 1269 (2021).

[44] J. F. Pelletier, L. Sun, K. S. Wise, N. Assad-Garcia, B. J. Karas, T. J. Deerinck, M. H. Ellisman, A. Mershin, N. Gershenfeld, R.-Y. Chuang, J. I. Glass, and E. A. Strychalski, Genetic requirements for cell division in a genomically minimal cell, Cell 184, 2430 (2021).

[45] A. B. O. Junior, V. G. Contessoto, M. F. Mello, and J. N. Onuchic, A scalable computational approach for simulating complexes of multiple chromosomes, Journal of molecular biology 433, 10.1016/J.JMB.2020.10.034 (2021).

[46] P. Eastman, J. Swails, J. D. Chodera, R. T. McGibbon, Y. Zhao, K. A. Beauchamp, L.-P. Wang, A. C. Simmonett, M. P. Harrigan, C. D. Stern, R. P. Wiewiora, B. R. Brooks, and V. S. Pande., Openmm 7: Rapid development of high performance algorithms for molecular dynamics., PLOS Comp. Biol. 13, e1005659 (2017).

[47] W. Humphrey, A. Dalke, and K. Schulten, VMD – Visual Molecular Dynamics, Journal of Molecular Graphics 14, 33 (1996).

